# Multiscale modelling of chromatin 4D organization in SARS-CoV-2 infected cells

**DOI:** 10.1101/2023.07.27.550709

**Authors:** Andrea M. Chiariello, Alex Abraham, Simona Bianco, Andrea Esposito, Francesca Vercellone, Mattia Conte, Andrea Fontana, Mario Nicodemi

## Abstract

SARS-CoV-2 is able to re-structure chromatin organization and alters the epigenomic landscape of the host genome, though the mechanisms that produce such changes are still poorly understood. Here, we investigate with polymer physics chromatin re-organization of the host genome, in space and time upon SARS-CoV-2 viral infection. We show that re-structuring of A/B compartments is well explained by a re-modulation of intra-compartment homotypic affinities, which leads to the weakening of A-A interactions and enhances A-B mixing. At TAD level, re-arrangements are physically described by a general reduction of the loop extrusion activity coupled with an alteration of chromatin phase-separation properties, resulting in more intermingling between different TADs and spread in space of TADs themselves. In addition, the architecture of loci relevant to the antiviral interferon (IFN) response, such as DDX58 or IFIT, results more variable within the 3D single-molecule population of the infected model, suggesting that viral infection leads to a loss of chromatin structural specificity. Analysis of time trajectories of pairwise gene-enhancer and higher-order contacts reveals that such variability derives from a more fluctuating dynamics in infected case, suggesting that SARS-CoV-2 alters gene regulation by impacting the stability of the contact network in time.

Overall, our study provides the first polymer-physics based 4D reconstruction of SARS-CoV-2 infected genome with mechanistic insights on the consequent gene mis-regulation.

## Introduction

SARS-CoV-2 outbreak had huge impact on society and science. Several efforts have been done to understand the dramatic effects of the virus action on host cell from different point of views, ranging from the immunological response^1^ to the effects of the infection on the epigenetic regulation^2^ or research of therapeutic molecular targets^3^. On the other hand, SARS-CoV-2 is able to impact chromatin architecture^4,5^ of the host cell, which in general is an important control layer for gene regulation^6,7^. Indeed, virus infection has been shown, for instance, to alter genome organization of olfactory receptors in humans and hamsters, providing a potential mechanism to explain anosmia^5^, one of the typical symptoms of Covid-19. More recently, it has been discovered that SARS-CoV-2 deeply impacts genome organization at multiple length scales, influencing the activity of gene categories^4^ fundamental to immunological response^1^, as interferon (INF) response and pro-inflammatory genes. Although those studies shed light on SARS-CoV-2 effects on genome organization, the physical mechanisms regulating how the virus changes the host cell 3D chromatin structure are not clearly understood.

Here, we employ models from polymer physics^8,9^ and Molecular Dynamics (MD) simulations to quantitatively study multiscale chromatin re-arrangements resulting from SARS-CoV-2 infection of the host cell. At large genomic length scales (several Mb), we show that a simple block-copolymer model^10,11^, based on combination of homo- and hetero-typic interactions, is able to explain the experimentally observed weakening of A compartment and enhancement of A-B mixing^4^, by basically reducing the intra-compartment A-A interaction affinity. At TAD level (from hundreds Kbs to some Mbs), we show that a model combining loop-extrusion^12,13^ and phase-separation^11,14^ effectively describes the experimentally observed intra-TAD weakening in SARS-CoV-2 infected cells^4^, by reducing the density of extruders coupled with an alteration of phase-separation properties of chromatin filament. Furthermore, using the same model informed with HiC data^15,16^, we investigate the architecture of genomic loci containing DDX58 and IFIT genes, which are of relevant immunological interest since linked to the antiviral interferon (IFN) response^17^ of the host cell. Specifically, analysis of polymer structures reveals that in SARS-CoV-2 model the population of single-molecule 3D configurations results more variable with respect to non-infected condition, suggesting that the alteration of activity observed for INF genes^1,4^ can be due to a general loss of structural specificity. By leveraging on our Molecular Dynamics simulations, we show that the model of SARS-CoV-2 exhibits a more scattered time dynamics, leading to a reduction of contact stability between pairs or hubs of multiple regulatory elements.

Overall, our polymer-physics based study provide mechanistic insights on how viral infection affects chromatin organization and provides a potential link between the observed genome re-structuring and gene mis-regulation within the host cell.

## Results

We study chromatin re-organization of host cell genome infected by SARS-CoV-2. To this aim, we consider recently published HiC data^4^ in control condition, i.e. A549 mock infected cells (referred to as Mock) and in A549 cells at 24-hour post SARS-CoV-2 infection, in which HiC data highlighted re-arrangements at multiple length scales, involving A/B compartment, TADs and regulatory contacts within specific loci^4^.

### Modelling of chromatin re-structuring in A/B compartments

One of the main structural re-arrangements on chromatin architecture resulting from SARS-CoV-2 infection of the host genome occurs at A/B compartment level. Specifically, it has been observed that viral infection results in a general weakening of A-compartment concomitantly with an enhanced A/B compartment mixing^4^, as schematically depicted in Figure 1A. To quantitatively investigate such effect, we first focused on a simple model of chromatin at A/B compartment level. We employed the Strings and Binders Switch polymer model^11,18^, where chromatin folding is driven by a phase-separation mechanism (Methods), similar to other models proposed for chromatin compartmentalization^10,19–21^. Briefly, we consider a simple block co-polymer where A and B compartments are modelled as two different types of binding sites (represented as different colors) which can homo-typically interact with cognate molecules (named binders) with an affinity E_A-A_ and E_B-B_, driving A-A or B-B interactions within the same compartment (Figure 1B). On the other hand, binders can also mediate A-B or B-A heterotypic interactions, with a general affinity E_A-B_ (Methods). To ensure micro-phase separation of A and B blocks, we always consider E_A-A_>E_A-B_ and E_B-B_>E_A-B19_. We first considered models with balanced interactions E_A-A_=E_B-B_ and varied the homotypic affinity (for sake of simplicity, heterotypic affinity E_A-B_ is kept constant, Methods). In general, low homo-typic interactions result in a reduced compartmentalization and increased A/B mixing, as shown by the model contact maps (Figure 1C and Suppl. Fig. 1A), the first eigenvector E1 from PC analysis and the saddle-plots of the sorted eigenvector components^22,23^ (Suppl. Fig. 1B, Methods). In addition, models with unbalanced interactions with E_B-B_>E_A-A_ result in both A/B mixing and, importantly, in weakening of A-compartment shown in the contact maps (Figure 3B, Suppl. Fig. 1C) and in asymmetric saddle plots (Suppl. Fig. 1D). Therefore, we reasoned that a combination of models with balanced and unbalanced interactions can fit A/B compartment alteration in SARS-CoV-2 infected genomes. Therefore, we fitted the best combination of interactions to reproduce the average compartment profile (using saddle-plot maps) obtained from HiC data in Mock and SARS-CoV-2-infected cells (Methods). Interestingly, Mock HiC data are mainly described (almost 90%) by a model with balanced homotypic interactions (i.e. E_A-A_=E_B-B_ Figure 1D, bottom left panel), indicating a substantial similarity in the A and B compartmentalization level and consistent with existing models of A/B compartmentalization^20^. Conversely, data in SARS-CoV-2 infected cells are best described by a combination of unbalanced homotypic interactions where E_B-B_>E_A-A_ (>60%) consistently with the general weakening of A-compartment and above-mentioned enhanced A/B mixing, with balanced interactions only marginally involved (about 20%, Figure 1D, bottom right). Importantly, albeit very simple, this model exhibits a high level of agreement with experimental data, as shown by the comparison between Log2 FC (SARS-CoV-2/Mock) of saddle-plot matrices (Pearson r=0.77, Figure 1E). Overall, these results show that chromatin re-arrangements observed in infected host genome can be explained by a re-modulation of affinities which in turn affect the tendency of compartments to microphase separate, as also shown by the 3D rendering of polymer structures representing A and B compartments in Mock (Figure 1F, left panel) and SARS-CoV-2 (Figure 1F, right panel) infected conditions.

**Figure 1:**
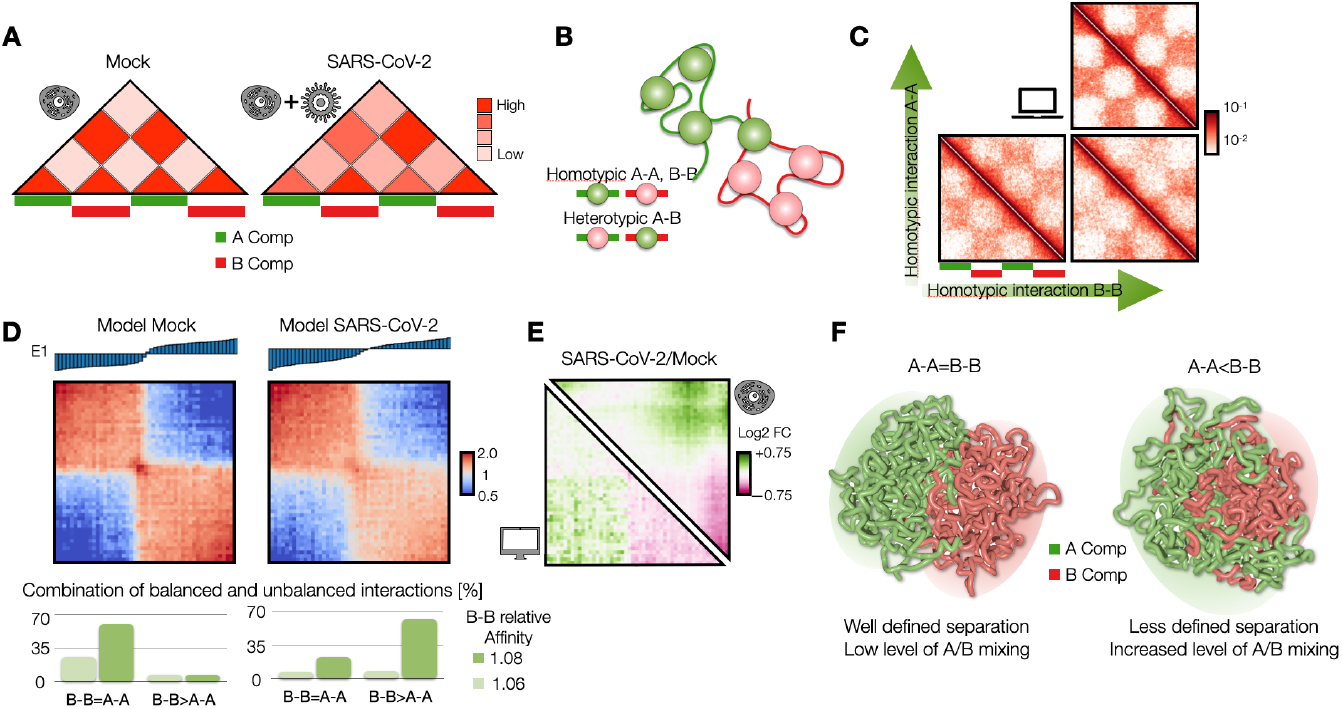
Modelling of chromatin re-structuring in A/B compartments. **A**: A/B compartment mixing in SARS-CoV-2 infected genome detected from HiC data^4^. **B**: Polymer model of A/B compartment envisages homotypic (A-A and B-B) and heterotypic (A-B) interactions. **C**: Variation of homotypic binding affinity results in weakening of A-compartment and enhancement of A/B compartment mixing. Heatmaps are computed from a populations of 3D structures obtained from MD simulations. **D**: Saddle-plot of best fit polymer model for Mock and SARS-CoV-2 conditions. Above, the sorted 1^st^ eigenvector E1 is shown. Below, best fit coefficients obtained to fit experimental saddle-plots. Dark and light green bars indicate B-B affinity used in the fit, normalized with respect the background heterotypic interaction (Methods). **E**: Log2 fold change of the saddle-plots from the model (bottom left) and HiC data^4^ (top right). Pearson correlation between matrices is r=0.77. **F**: 3D rendering of best polymer models for Mock (left) and SARS-CoV-2 (right).

### Viral infection impacts loop-extrusion and phase-separation features at TAD level

Next, we investigated how SARS-CoV-2 infection impacts genome organization at TAD level, i.e. genomic scales ranging from tens of kbs to some Mbs. Indeed, it has been shown that viral infection produces a general weakening of intra-TAD contacts along with a slightly increase of inter-TADs interactions^4^ (Figure 2A) and concomitantly with a general reduction of Cohesin level^4^, suggesting a reduction of loop extrusion activity. To test this hypothesis and give a mechanistic insight to this result, we used a polymer physics model combining both loop-extrusion^12,13^ (LE) and phase-separation^11,18^ (PS) mechanism (Figure 2B, Methods), which recently has been shown to successfully describe chromatin organization at single cell level^16^. In this scenario, LE and PS simultaneously act and the pattern of chromatin contacts observed in HiC data results from an interplay between both processes (Figure 2C). By varying the main system parameters, i.e. interaction affinity and average distance between extruders (or equivalently their number, Methods) (Suppl. Fig. 2A), we generated several different polymer populations with their simulated contact maps (Suppl. Fig. 2B) and contact probability profiles (Suppl. Fig. 2C). In this way, we were able to identify the polymer model best fitting the contact probability obtained from HiC data (Methods), in the genomic distance ranging from the sub-TAD level (approx. 10kb) to inter-TADs contacts (some Mbs, Methods). The model is able to explain with accuracy experimental data, as shown by the fit of the average contact probability (as shown by χ^2^ values in Suppl. Fig. 2D) in Mock (Figure 2D, left bottom panel) and in SARS-CoV-2 infected (Figure 2D, right bottom panel) conditions. Importantly, the best model describing Mock data revealed an average distance between extruders of approximately 100-150 kb (Suppl. Fig. 2D, left panel), consistent with previous estimates obtained from other HiC datasets^13^. Conversely, the best model fitting the SARS-CoV-2 infected HiC data was best described by a consistently decreased number of extruders (approximately halved, Suppl. Fig. 2D, right panel), in full agreement with experimental observations where viral infection produces a genome-wide decrease of Cohesin levels^4^. Interestingly, this analysis revealed that, in order to fit HiC data in infected cells, the reduction of extruders is coupled with a reduction of interaction affinity (around 15-20%) between binders and chromatin (Suppl. Fig. 2D), which affects chromatin spatial localization and contributes to the general weakening of intra-TADs contacts (Figure 2D, upper panels) observed in infected genomes. This is quantitatively shown in the Log2 FC (SARS-CoV-2/Mock) of contact maps (Figure 2E, upper panel) and contact probabilities (Figure 2E, bottom panel), which exhibits a very good agreement with experimental data (Pearson r=0.82, Methods). To check the biological robustness of our results, we repeated the above-described analysis to HiC data in cells infected by the human coronavirus (HCoV) OC43^4^, which causes common cold, used as control case. Again, we were able to fit the average contact probability with high accuracy (chi-square test p-val=1). Intriguingly, we find that the best model is mainly analogous to the Mock case, with same affinity and a light increase of average distance between extruders (as observed in SARS-CoV-2, but to a lesser extent), as shown by the best fitting parameters (Suppl. Fig. 2E) and in full agreement with the experimental reports^4^. Finally, taking advantage of MD simulations, we produced an example of 3D structure representing the average TAD in Mock (Figure 2F, left panel and Suppl. Video 1) and SARS-CoV-2 infected (Figure 2F, right panel and Suppl. Video 2) conditions, providing an effective and realistic summary of the architectural re-arrangements occurring within and between TADs after the infection. Overall, our simulations suggest that viral infection affects genome organization by altering fundamental physical mechanisms, including loop-extrusion and phase-separation, that shape chromatin structure.

**Figure 2:**
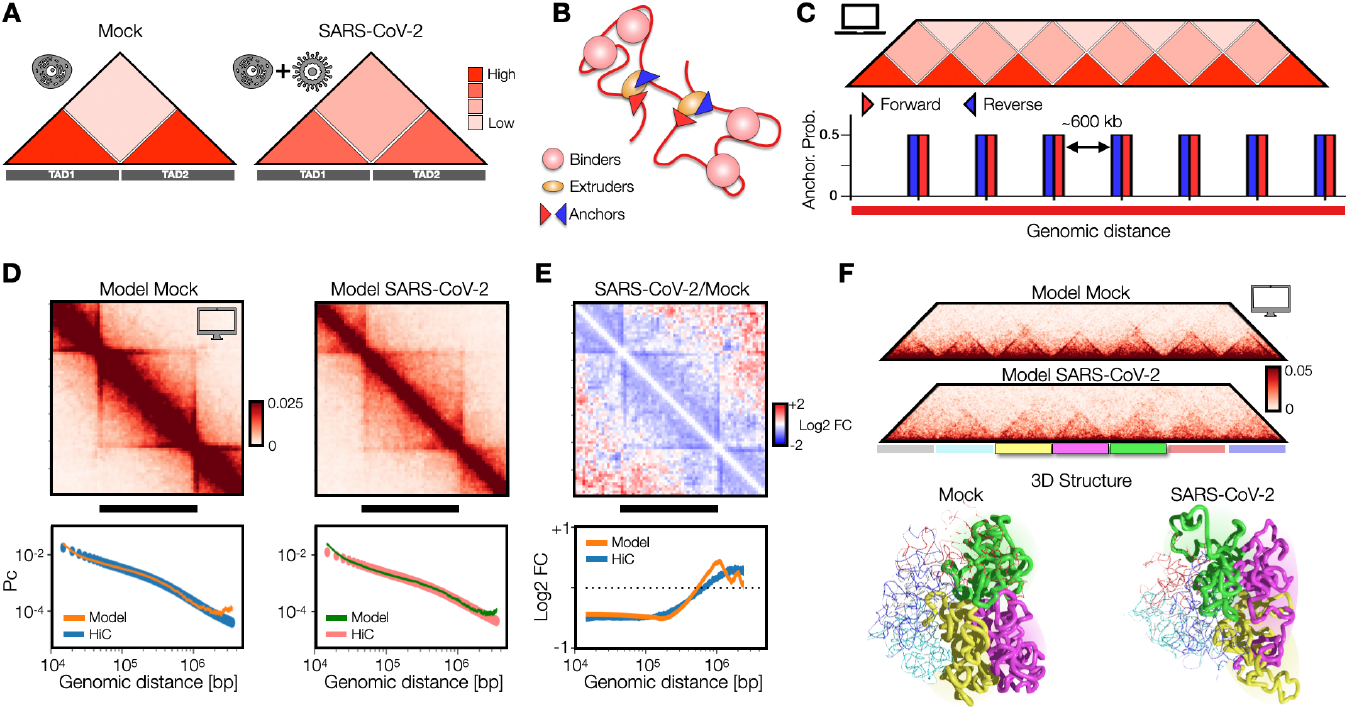
Viral infection impacts loop-extrusion and phase-separation features at TAD level. **A**: Intra-TAD weakening in SARS-CoV-2 infected genome detected from HiC data^4^. **B**: Polymer model of genome organization at TAD scale envisages chromatin loop-extrusion and interactions between chromatin and proteins (binders). **C**: TAD boundaries are limited by converging (forward-reverse) anchor points, occurring with a probability (Methods). **D**: Average contact maps of TADs from the best model for Mock (left) and SARS-CoV-2 conditions, obtained fitting experimental contact probabilities (Methods). Below, best fit and experimental contact probabilities. **E**: Log2 Fold Change of average contact maps. Below, Log2 Fold Change of contact probabilities in HiC^4^ (blue curve) and model (orange curve). Correlation between the curves is r=0.82. **F**: Contact maps of best fit models for Mock and SARS-CoV-2 conditions. Below, 3D rendering of polymer structures obtained from MD simulations of Mock and SARS-CoV-2 models. For visualization purposes, only the three central TADs are shown.

### Structural re-arrangements of interferon response genes (IFN) loci

Next, to understand how the above discussed structural re-arrangement within TADs may affect gene regulation, we modelled real genomic regions relevant in case a viral infection occurs. Specifically, we considered genomic loci containing interferon (IFN) response genes, i.e. genes typically upregulated upon interferon stimulus and that are commonly expressed as response to a viral infection^17^. Importantly, it has been shown that in severe Covid syndromes such genes are not properly expressed^1,24^ with consequent alteration in the immunological response of host cell. We considered as first case of study the genomic region spanning 400kb around the DDX58 gene (chr9: 32300000-32700000 bp, hg19 assembly, Figure 3). The DDX58 locus exhibits the typical re-arrangements caused by SARS-CoV-2 infection, as in Mock case the DDX58 gene is contained in a well-defined domain limited by convergent CTCF sites (Figure 3A), whereas in the infected case a general weakening of intra-TAD interactions is observed, although CTCF peaks are mainly unchanged (Figure 3B). Analogous observations hold for another IFN locus, containing the cluster of IFIT genes (chr10: 90900000-91290000 bp) (Suppl. Fig. 3A and 3B). To quantitatively investigate such re-arrangements, we employed the above-described polymer model combining loop-extrusion and chromatin-protein interactions^16^, using experimental CTCF ChIP-seq data^4^ to set the probabilities and the positions of the anchor points for extruders^13^ and HiC data to optimize the types and the positions of the binding sites^15^ (Suppl. Fig. 4A, Methods). Taking advantage of the results obtained for the polymer model calibrated to simulate the average chromatin behavior at TAD level, we were able to generate ensembles of 3D structures accurately capturing the differences in the DDX58 locus between Mock and SARS-CoV-2 conditions, as shown by the simulated contact maps (Figure 3C and 3D) highly correlated with experimental data (Pearson r>0.9, distance corrected r’=0.67, Methods). In addition, the model correctly captures the different contact probability decay (Suppl. Fig. 4B), as shown by the Log2 FC curve (Suppl. Fig. 4C, Pearson r=0.81). Analogous results were found for the polymer model of the IFIT locus, which returns highly correlated contact maps (Suppl. Fig. 3C and 3D) and similar contact probability decays (Suppl. Fig. 4D and 4E). Finally, examples of 3D structures taken from MD simulations visually highlight the above-discussed architectural differences, with the DDX58 and IFIT loci organized in distinct, well-defined regions in Mock (Figure 3E and Suppl. Fig. 3E) while they tend to be less localized and more intermingled in SARS-CoV-2 (Figure 3F and Suppl. Fig. 3F).

**Figure 3:**
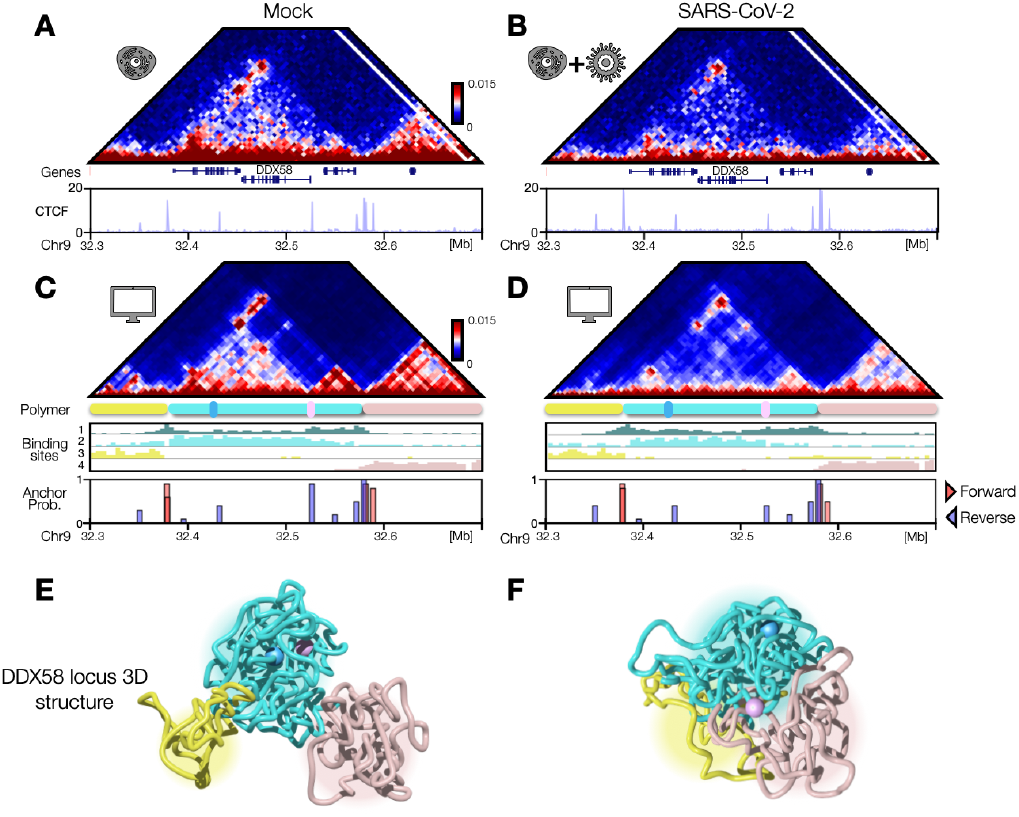
Structural re-arrangements of IFN DDX58 locus. **A**: Mock HiC data of the genomic region (chr9:32300000-32700000, hg19) centered around the interferon response DDX58 gene. Below, CTCF signal is shown (data taken from Ref.^4^). **B**: As panel A, SARS-CoV-2 HiC and CTCF data. **C**: Simulated contact map for Mock polymer model. Below, binding sites profile, anchor point probability and their orientation (Methods) is shown. **D**: As panel C, SARS-CoV-2 model. **E**: Example of 3D structure of DDX58 locus taken from an MD simulation of the Mock model. Different regions are differently colored according to the pattern of the contact maps. Pink and cyan spheres highlight the position of DDX58 and its enhancer respectively. **F**: As panel E, SARS-CoV-2 model.

### Single cell 3D structures result highly variable in SARS-CoV-2 infected condition

The different 3D structures observed in Mock and SARS-CoV-2 prompted us to investigate in more detail the above-discussed architectural differences at the single cell level. To this aim, polymer models offer a powerful tool as they allow to build ensembles of independent 3D structures that mimic single-cell variability^14^, experimentally observed e.g. by MERFISH microscopy method^25^. Therefore, leveraging on such feature, we analyzed the population of 3D structures in Mock (Figure 4A, upper panel) and SARS-CoV-2 (Figure 4A, bottom panel) models. First, we focused on the DDX58 promoter and its validated enhancer^4^ (Figure 4A).

**Figure 4:**
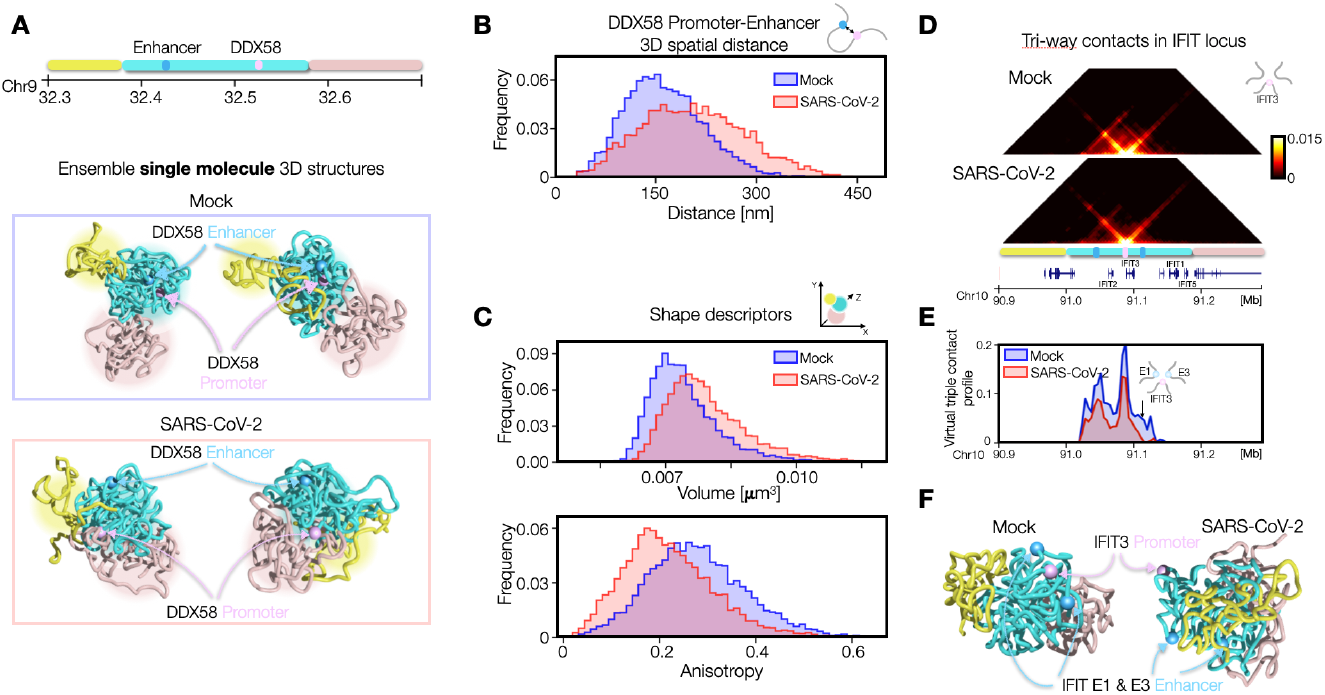
Single cell 3D structures result more variable in SARS-CoV-2 infected condition. **A**: Examples of 3D structures from the ensemble of single molecule configurations for DDX58 locus in Mock (top panel) and SARS-CoV-2 (bottom panel) models. DDX58 promoter and its enhancer are highlighted in pink and cyan respectively. **B**: Distributions of 3D distances between DDX58 promoter and its enhancer. Length scales are estimated by mapping the model in physical units (Methods). Distributions are statistically different (one-sided T-test, p-val=10^−259^). **C**: Size and shape descriptors computed from the entire polymer representing the DDX58 locus. Mock and SARS-CoV-2 models exhibit different volume (top panel, one-sided T-test p-val=10^−240^) and anisotropy (bottom panel, one-sided T-test p-val=10^−240^) distributions. **D**: Simulated matrices of triple contacts using IFIT3 as point of view in Mock (top panel) and SARS-CoV-2 (bottom panel) models. **E**: Virtual triple contact profile fixing enhancer 1 and IFIT3 as point of views. Arrow indicates probability of E1-IFIT3-E3 triple. **F**: Examples of 3D structures from the ensemble of single molecule configurations for IFIT locus in Mock (left) and SARS-CoV-2 (right) models. Positions of IFIT3 and enhancers E1 and E3 are highlighted in pink and cyan respectively.

By visual inspection of these 3D structures in both conditions, it emerges that DDX58 promoter and the enhancer tend to be closer in space in Mock with respect the infected condition, in agreement with HiC data. The distributions of 3D distances between the DDX58 promoter and its enhancer (Figure 4B) confirmed this observation, as in Mock it exhibits a lower mean than the infected case (one-sided T-test p-val=10^−259^). Interestingly, the distribution results also more variable in infected cells (st. dev. in SARS-CoV-2 ∼30% higher than in Mock), suggesting that the mis-regulation of this gene upon infection is also due to a loss of contact specificity in the population of 3D structures. Next, we focused on the architecture of the entire locus and considered the polymer size and shape descriptors ^26^ (Methods). Again, we find that the estimated volume distribution (Methods) is more variable in SARS-CoV-2 (Figure 4C upper panel, st. dev. ∼30% higher). Conversely, the average anisotropy distribution (Figure 4C, bottom panel), which measures how asymmetrically the polymer is distributed in space, results lower in SARS-CoV-2 population. Analogous results are found for a-sphericity, another shape descriptor (Methods) measuring the deviation from a spherical geometry. Those results are consistent with the increased inter-TADs contacts and less localization observed which make the polymer more homogeneous and spherical in SARS-CoV-2 model.

Next, we investigated whether the infected model may exhibit differences on higher-order contacts. To this aim, we focused on the cluster of IFIT genes, where we considered the probability of three-way contacts^27,28^ using as point of view IFIT3 gene, located in the center of the IFIT TAD (Figure 4D, Methods). We find that in SARS-CoV-2 model three-way contacts result consistently reduced (Figure 4D) although weak, long-range events appear. By fixing the enhancer 1 (E1) as other point of view we generated a virtual three-way profile involving E1 and IFIT3 (Figure 4E), which clearly highlights that specific three-way contacts are reduced in SARS-CoV-2 model. This suggests that the mis-regulation may also be due to an alteration of contact network within the regulatory hub, consistent with other recent observations whereby the olfactory hubs are disrupted/perturbed after SARS-CoV-2 infection^5^. Finally, examples of 3D structures of the IFIT locus in Mock (Figure 4F, left panel) and SARS-CoV-2 (Figure 4F, right panel) conditions provide a visual summary of the discussed results.

### Time dynamics of 3D contacts is highly variable in SARS-CoV-2

Next, we investigated the mechanism leading to the different structural variability observed in Mock and SARS-CoV-2 models. To this aim, we considered the population of independent time trajectories (Methods) of the polymer and analyzed the dynamics in both conditions (Figure 5A) at equilibrium (Methods). We focused again on the DDX58-enhancer distance (Figure 5B) and the above-discussed polymer shape descriptors, i.e. anisotropy (Suppl. Fig. 5A, left panel) and a-sphericity (Suppl. Fig. 5A, right panel). As expected, the distance trajectories in the Mock model appear fluctuating around average values lower than the SARS-CoV-2 model, as also confirmed by the distributions of the average distance over different time trajectories (Figure 5C, upper panel, T-test p-val=10^−15^, Methods). Same analysis for anisotropy (Suppl. Fig. 5B, left panel) and a-sphericity (Suppl. Fig. 5B, right panel) reveals instead a specular behavior, in agreement with the observations of the previous section. Interestingly, it emerges also that the time trajectories in SARS-CoV-2 model are more fluctuating, as shown by the distribution of the standard deviations of the distance in time (Figure 5C, lower panel). For the shape descriptors Mock and SARS-CoV-2 models exhibit similar deviations from the average value during time (Suppl. Fig. 5C). Analogously, multiple co-localization events (named co-occurrences, Methods) in IFIT locus, involving IFIT3 and two enhancers, tend to be less frequent in time in SARS-CoV-2 model dynamics (Suppl. Fig. 5D). These results suggest that SARS-CoV-2 could affect the stability of contacts between regulatory elements. To support this conclusion, we analyzed in more detail the DDX58-enhancer distance time dynamics by considering shorter time scales at higher time resolution (Figure 5D, Methods). We generated time trajectories to follow a smooth evolution of gene-enhancer distance, in Mock (Suppl. Video 3) and SARS-CoV-2 (Suppl. Video 4) models. In this way, we were able to estimate a “contact time”, i.e. how long the gene and the enhancer spend in contact (Figure 5E, Methods). Importantly, we find that the distribution of contact times tends to be significantly lower in SARS-CoV-2 model (Figure 5F, T-test p-val=2*10^−4^), with an approximately 1.5-fold reduction of the average time. Taken together, those results point toward a scenario where the mis-regulation of IFN genes observed in SARS-CoV-2 could be imputed to a decreased contact stability between genes and their regulatory elements.

**Figure 5:**
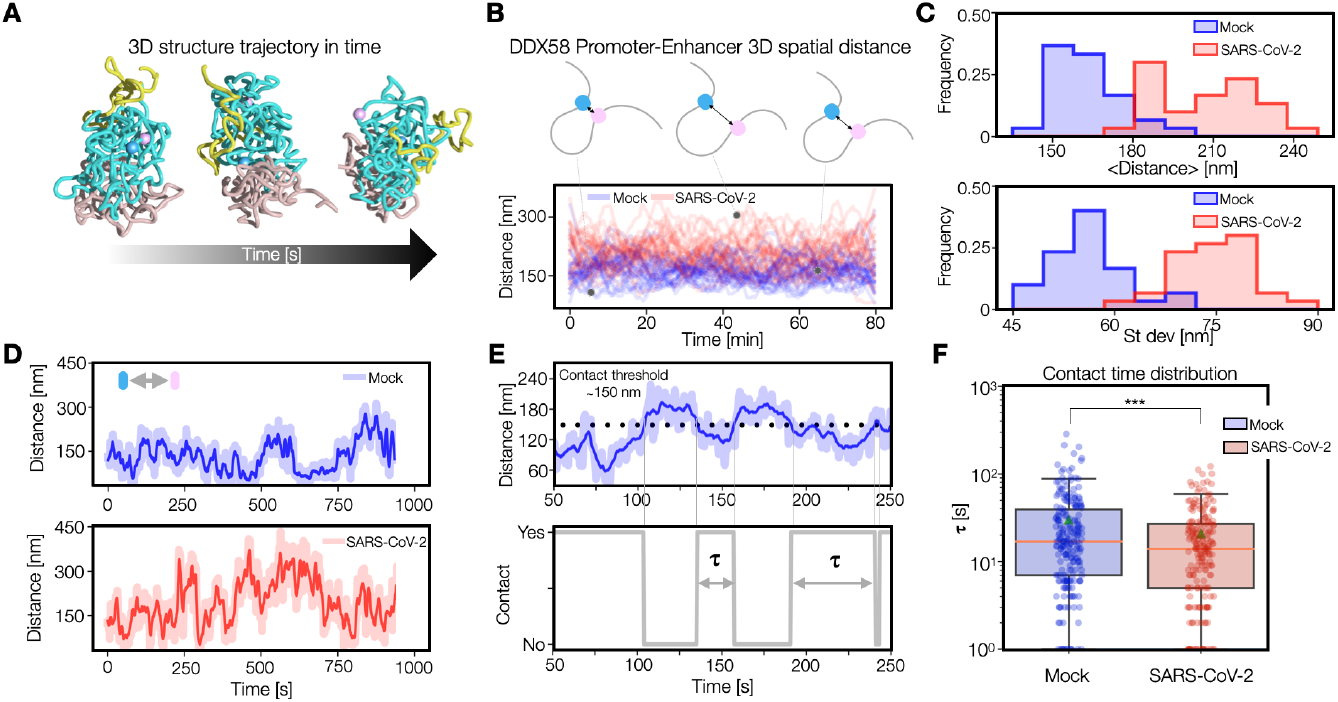
Time dynamics of 3D contacts is more variable in SARS-CoV-2. **A**: Evolution of 3D structure during time. **B**: 3D distance trajectories between DDX58 promoter and its enhancer in Mock (blue curves) and SARS-CoV-2 (red) models. **C**: Distributions of average distance (top) and standard deviation (bottom) computed over independent time trajectories shown in panel B. In both cases, Mock is statistically different from SARS-CoV-2 model (p-val=10^−15^ and 10^−18^ respectively, one-sided T-test). **D**: 3D distance trajectories at higher time resolution in Mock and SARS-CoV-2 models. Darker lines are smoothed curves (Methods). **E**: Given the 3D distance trajectory (top) we define the contact time **τ** (bottom) as the time period spent below the contact threshold, here set to 150nm (Methods). **F**: Boxplots showing the distribution of contact time **τ** in Mock (blue) and SARS-CoV-2 (red) models. Distributions have statistically different averages (p-val=2*10^−4^, one-sided T-test).

### Chromatin re-arrangements in SARS-CoV-2 infection correlate with a combination of changes of CTCF and histone marks

Next, to understand the link between the architectural re-arrangements encoded in HiC data and molecular factors, we investigated the relationship between binding sites and epigenetics marks, such as CTCF and histone modifications. To this aim, we made a cross-correlation analysis (Methods) between the binding site profiles of the model and different available epigenetic marks at DDX58 (Figure 6) and IFIT (Suppl. Fig. 6) loci, in Mock and SARS-CoV-2 conditions. In Mock, we find (Figure 6A, right panel) a clear, strong correlation between CTCF and RAD21 with binding site type #1, likely highlighting an important role for LE mechanism in shaping the central domain containing the DDX58 gene, but we also observe a significative correlation with RNAPolII (RPB1) and H3K4me3, in agreement with the view of a combinatorial action of different factors in shaping chromatin organization^29^. In addition, it emerges a clear association between the flanking binding sites (#3 and #4) to H3K27me3 and H3K9me3 respectively (Figure 6B, left panel). In SARS-CoV-2 model the distribution of binding sites exhibits, in general, a similar profile (Figure 6A, left panel) but a richer pattern of (less strong) correlations is found (Figure 6B, right panel), as CTCF and RAD21 (generally reduced) are associated with multiple types (#1 and #3) and changes in correlations involving H3K27ac, which exhibits a general reduction too^4^, occur. Analogous considerations hold for IFIT locus where changes in correlations involve CTCF, RPB1 and H3K4me3 (Suppl. Fig. 6). In general, those results support the proposed mechanism^4^ by which an alteration of LE activity upon infection coupled with changes in the epigenetic signatures of activity produces an altered expression of INF genes with a consequent poor response to the infection.

**Figure 6:**
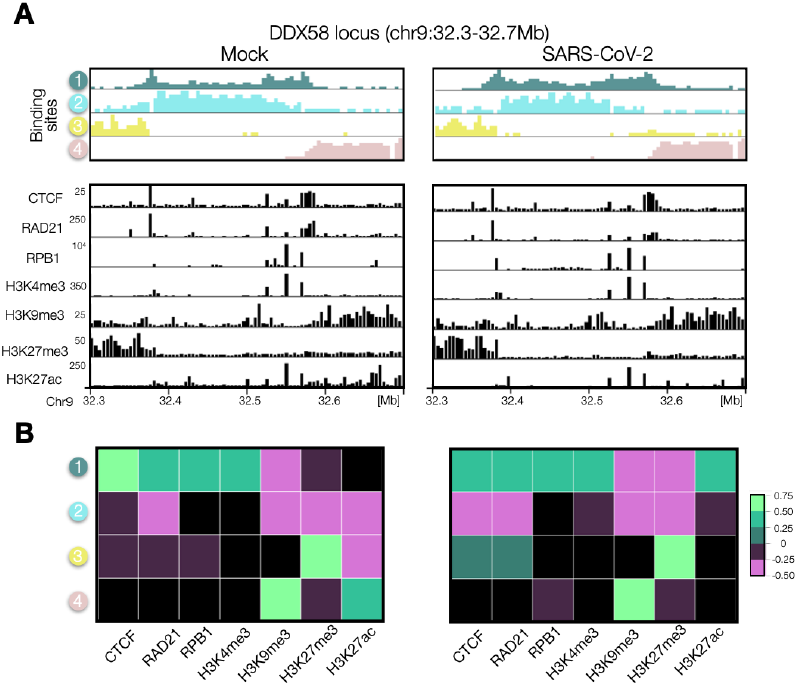
Chromatin re-arrangements correlates with a combination of changes of CTCF and histone marks at DDX58 locus. **A**: Top panel: distribution of binding sites obtained from HiC data in Mock (left) and SARS-CoV-2 (right) conditions. Bottom panel: different epigenetic marks (data from Ref.^4^). In SARS-CoV-2 relevant reductions of RAD21 and H3K27ac are observed. **B**: Cross-correlation analysis between binding site profiles and epigenetic marks. Significative correlations (Methods) at DDX58 locus in Mock (left) and SARS-CoV-2 (right) models are shown.

## Discussion

In this work, we investigated how SARS-CoV-2 infection alters the 3D organization of chromatin in the host cell at multiple length scales, ranging from few kilobases to several Mbs. To this aim, we employed models from polymer physics and MD simulations widely used to study chromatin organization^11,18,30^. We showed that a simple block-polymer including just homo-typic and hetero-typic interactions is overall able to describe the observed A-compartment weakening and A-B mixing, by remodulating A-A affinities in an unbalanced A/B compartment model. Of course, more complicated descriptions of compartmentalization are possible and could include other mechanisms known to play a role for chromatin structure, such as interaction with nuclear envelope^19,31^. At TAD level, we find that a combined reduction of loop extrusion activity (modelled as a reduction of extruders) together with an alteration of phase-separation properties (modelled as a reduction of chromatin-protein affinities) potentially explain the weakening of intra-TAD interactions observed in HiC data^4^. Interestingly, a control model calibrated from HiC data in host cells infected with common cold virus^4^ has a slightly reduced loop-extrusion activity with respect the Mock case but keeps unchanged protein affinities with chromatin, suggesting that the capacity of altering this phase-separation properties is a peculiar feature of SARS-CoV-2 model. We then investigated the link between chromatin re-arrangement and the regulation of genes involved in the antiviral response (IFN genes) which are mis-regulated upon SARS-CoV-2 infection^1^. Polymer models of genomic loci containing DDX58 and IFIT genes highlighted a higher degree of variability in the ensemble of single-molecule conformations of SARS-CoV-2 models. This variability is in turn related to a noisier and less stable time dynamics, suggesting that SARS-CoV-2 infection reduces specificity and structural stability of regulatory contacts.

In order to understand the molecular causes leading to the above-discussed re-arrangements, by means of direct or undirect mechanisms, it would be interesting to integrate in the polymer model the existence of specific molecular factors encoded by the virus known to perturb the host cell, as highlighted by recent experiments showing that viral proteins can alter the cell epigenome^2^ (ORF8) or interact with other proteins of the infected cell^32^. In this regard, it is worth to mention that other virus are capable of re-structuring genome organization through the transcription of their proteins, such as NS1 from influenza A virus (IAV)^33^. This strategy could be relevant to test the effects of specific proteins on the physical mechanisms shaping chromatin architecture (e.g. phase-separation) and therefore be helpful in the identification of molecular targets for therapeutics purposes.

In general, exploring the link between viral infection and chromatin architecture can be extremely insightful to understand virus action on host cell at the level of gene regulation. To this aim, polymer models turn out to be valuable tool as they offer an unbiased, predictive approach to connect different aspects relevant for genome organization and function^34^, including single-cell variability, dynamics between regulatory elements and research of therapeutic targets.

## Acknowledgements

A.M.C. acknowledges “Programma per il Finanziamento della Ricerca di Ateneo Linea B” (FRA) 2020, University of Naples Federico II, CINECA ISCRA Grant ID PhaSSep - HP10C8JWU7. M.N. acknowledges support from the National Institutes of Health Common Fund 4D Nucleome Program grant 5 1UM1HG011585-03, EU H2020 Marie Curie ITN n.813282, PNRR MUR M4C2 CN00000041 “National Center for Gene Therapy and Drugs based on RNA Technology” NextGenerationEU CUP E63C22000940007, MUR PRIN 2022 2022R8YXMR, and computer resources from INFN, CINECA, ENEA CRESCO/ ENEAGRID88 and Ibisco at the University of Naples.

## Author contributions

A.M.C and M.N. designed the project. A.M.C. and A.A. developed the modeling part; A.M.C., A.A. S.B., A.E. ran the computer simulations and performed their analyses. A.M.C. wrote the manuscript with input from the other authors.

## References

1. Carvalho, T., Krammer, F. & Iwasaki, A. The first 12 months of COVID-19: a timeline of immunological insights. Nat. Rev. Immunol. 2021 214 21, 245–256 (2021).

2. Kee, J. et al. SARS-CoV-2 disrupts host epigenetic regulation via histone mimicry. Nat. 2022 6107931 610, 381–388 (2022).

3. Ho, J. S. Y. et al. TOP1 inhibition therapy protects against SARS-CoV-2-induced lethal inflammation. Cell 184, 2618-2632.e17 (2021).

4. Wang, R. et al. SARS-CoV-2 restructures host chromatin architecture. Nat. Microbiol. 2023 84 8, 679–694 (2023).

5. Zazhytska, M. et al. Non-cell-autonomous disruption of nuclear architecture as a potential cause of COVID-19-induced anosmia. Cell 185, 1052-1064.e12 (2022).

6. Kempfer, R. & Pombo, A. Methods for mapping 3D chromosome architecture. Nat. Rev. Genet. 21, 207–226 (2020).

7. Misteli, T. The Self-Organizing Genome: Principles of Genome Architecture and Function. Cell 183, 28–45 (2020).

8. Bianco, S. et al. Computational approaches from polymer physics to investigate chromatin folding. Curr. Opin. Cell Biol. 64, (2020).

9. Brackey, C. A., Marenduzzo, D. & Gilbert, N. Mechanistic modeling of chromatin folding to understand function. Nat. Methods 2020 178 17, 767–775 (2020).

10. Jost, D., Carrivain, P., Cavalli, G. & Vaillant, C. Modeling epigenome folding: Formation and dynamics of topologically associated chromatin domains. Nucleic Acids Res. 42, 9553–9561 (2014).

11. Chiariello, A. M., Annunziatella, C., Bianco, S., Esposito, A. & Nicodemi, M. Polymer physics of chromosome large-scale 3D organisation. Sci. Rep. 6, 29775 (2016).

12. Sanborn, A. L. et al. Chromatin extrusion explains key features of loop and domain formation in wild-type and engineered genomes. Proc. Natl. Acad. Sci. 112, E6456–E6465 (2015).

13. Fudenberg, G. et al. Formation of Chromosomal Domains by Loop Extrusion. Cell Rep. 15, 2038–2049 (2016).

14. Conte, M. et al. Polymer physics indicates chromatin folding variability across single-cells results from state degeneracy in phase separation. Nat. Commun. 11, 3289 (2020).

15. Bianco, S. et al. Polymer physics predicts the effects of structural variants on chromatin architecture. Nat. Genet. 50, 662–667 (2018).

16. Conte, M. et al. Loop-extrusion and polymer phase-separation can co-exist at the single-molecule level to shape chromatin folding. Nat. Commun. 2022 131 13, 1–13 (2022).

17. Schneider, W. M., Chevillotte, M. D. & Rice, C. M. Interferon-Stimulated Genes: A Complex Web of Host Defenses. https://doi.org/10.1146/annurev-immunol-032713-120231 32, 513–545 (2014).

18. Barbieri, M. et al. Complexity of chromatin folding is captured by the strings and binders switch model. Proc. Natl. Acad. Sci. 109, 16173–16178 (2012).

19. Falk, M. et al. Heterochromatin drives compartmentalization of inverted and conventional nuclei. Nat. 2019 5707761 570, 395–399 (2019).

20. Shi, G., Liu, L., Hyeon, C. & Thirumalai, D. Interphase human chromosome exhibits out of equilibrium glassy dynamics. Nat. Commun. 9, 3161 (2018).

21. Shin, S., Shi, G. & Thirumalai, D. A method for extracting effective interactions from Hi-C data with applications to interphase chromosomes and inverted nuclei.

22. Abdennur, N. & Mirny, L. A. Cooler: scalable storage for Hi-C data and other genomically labeled arrays. Bioinformatics 36, 311–316 (2020).

23. Open2C et al. Cooltools: enabling high-resolution Hi-C analysis in Python. bioRxiv 2022.10.31.514564 (2022) doi:10.1101/2022.10.31.514564.

24. Blanco-Melo, D. et al. Imbalanced Host Response to SARS-CoV-2 Drives Development of COVID-19. Cell 181, 1036-1045.e9 (2020).

25. Bintu, B. et al. Super-resolution chromatin tracing reveals domains and cooperative interactions in single cells. Science (80-.). 362, eaau1783 (2018).

26. Arkin, H. & Janke, W. Gyration tensor based analysis of the shapes of polymer chains in an attractive spherical cage. J. Chem. Phys. 138, (2013).

27. Chiariello, A. M. et al. A Dynamic Folded Hairpin Conformation Is Associated with α-Globin Activation in Erythroid Cells. Cell Rep. 30, 2125-2135.e5 (2020).

28. Oudelaar, A. M. et al. Single-allele chromatin interactions identify regulatory hubs in dynamic compartmentalized domains. Nat. Genet. 50, 1744–1751 (2018).

29. Esposito, A. et al. Polymer physics reveals a combinatorial code linking 3D chromatin architecture to 1D chromatin states. Cell Rep. 38, 110601 (2022).

30. Brackley, C. A., Taylor, S., Papantonis, A., Cook, P. R. & Marenduzzo, D. Nonspecific bridging-induced attraction drives clustering of DNA-binding proteins and genome organization. Proc. Natl. Acad. Sci. U. S. A. 110, E3605–E3611 (2013).

31. Ringel, A. R. et al. Repression and 3D-restructuring resolves regulatory conflicts in evolutionarily rearranged genomes. Cell 185, 3689-3704.e21 (2022).

32. Gordon, D. E. et al. A SARS-CoV-2 protein interaction map reveals targets for drug repurposing. Nat. 2020 5837816 583, 459–468 (2020).

33. Heinz, S. et al. Transcription Elongation Can Affect Genome 3D Structure. Cell 174, 1522-1536.e22 (2018).

34. Dekker, J. et al. The 4D nucleome project. Nature vol. 549 219–226 (2017).

35. Lieberman-Aiden, E. et al. Comprehensive mapping of long-range interactions reveals folding principles of the human genome. Science (80-.). 326, 289–293 (2009).

36. Kremer, K. & Grest, G. S. Dynamics of entangled linear polymer melts: A molecular-dynamics simulation. J. Chem. Phys. 92, 5057–5086 (1990).

37. Allen, M. P. & Tildesley, D. J. Computer Simulation of Liquids (Oxford Science Publications) SE - Oxford science publications. Oxford Univ. Press (1989).

38. Anderson, J. A., Glaser, J. & Glotzer, S. C. HOOMD-blue: A Python package for high-performance molecular dynamics and hard particle Monte Carlo simulations. Comput. Mater. Sci. 173, 109363 (2020).

39. Buckle, A., Brackley, C. A., Boyle, S., Marenduzzo, D. & Gilbert, N. Polymer Simulations of Heteromorphic Chromatin Predict the 3D Folding of Complex Genomic Loci. Mol. Cell 72, 786-797.e11 (2018).

